# Postural demands modulate tactile perception in the lower limb in young and older adults

**DOI:** 10.1101/2024.08.28.610073

**Authors:** Fabian Dominik Wachsmann, Katja Fiehler, Dimitris Voudouris

## Abstract

Balance control requires constant integration of feedforward and feedback signals. In healthy aging, the quality of feedback signals decreases while feedforward control is upweighted; but it is unclear how tactile perception is modulated when balance control is challenged and how this interacts with age-related changes in sensorimotor processes. We therefore examined tactile perception in standing when confronted with different postural demands in young and older adults. To this end, we measured tactile sensitivity on the calf during sitting (baseline), standing on solid ground, and standing on unstable ground (foam). We also measured the center of pressure during standing using a force plate and calculated a 95% confidence ellipse area and the center of pressure length. Tactile sensitivity was assessed by fitting a psychometric function to verbal responses for detecting vibrotactile probes, calculating the detection threshold at 50% detection, and normalizing the two standing conditions to baseline. We examined the effect of age and postural demands on the center of pressure kinematics and detection thresholds. We found higher sway and poorer tactile sensitivity when standing on foam irrespective of age. The increase of postural demands seems to reduce the reliance on tactile feedback signals from the lower limbs in both young and older adults. Our results suggest that postural demands challenge healthy agers as young adults, probably leading to a down-weighting of tactile feedback processing.

## Introduction

Standing upright appears a rather easy task, however, it requires the continuous integration of feedback and feedforward signals to generate the appropriate motor commands. To achieve a stable stance, humans typically integrate visual, vestibular, proprioceptive, and tactile signals based on their reliability [1,2], and combine these signals with sensorimotor predictions about future sensory states based on descending motor commands [3]. Consequently, postural control can deteriorate when sensory signals are impaired [2].

Sensorimotor predictions can lead to suppression of associated sensory feedback, a phenomenon called sensory suppression. In touch, this is reflected in reduced tactile sensitivity on the moving limb, as has been shown for various movements, from reaching and grasping [4,5] to standing [6]. Tactile suppression is explained by central efference copy mechanisms [7–9], though peripheral postdictive processes may also play some role [10,11]. The strength of tactile suppression can be modulated by multiple factors. For example, tactile suppression is weaker on the stationary leg that supports gait initiation [14]. Contrary, tactile suppression is stronger when task demands are high [15,but see also 16], probably by down-weighing task-irrelevant somatosensory feedback signals.

Healthy aging is associated with poorer and more delayed sensorimotor feedback control [17,18]. For instance, tactile signals from the foot sole are impaired which may lead to increased postural sway [19–21]. However, there are also reports that postural sway can be lower in older compared to young women [23]. Meanwhile, reliance on sensorimotor predictions may change with aging. For example, anticipatory postural adjustments before self-imposed perturbations are both smaller and delayed in older than young adults [24], though this can improve with suitable training [25]. Moreover, tactile suppression as an index of sensorimotor predictions is stronger in older adults [26,27,but see also 28], which might reflect a compensatory mechanism to adapt to their poorer sensory processing by relying more on predictive control. So far, it is unknown how tactile perception on the leg, which is elementary for postural control, is modulated during stance retention under different postural demands and how this is affected by old age that comes with impaired tactile perception.

Here, we examined how tactile perception is modulated by postural demands in young and older adults. We expected increased postural sway with higher postural demands in both age groups. Additionally, postural sway may be higher [20] or lower [23] in aging. Concerning tactile perception, we expected generally lower tactile sensitivity in older than young adults [17,26]. As sensorimotor feedback control is impaired in aging [e.g.18], older adults may rely more on predictive control [e.g. 27,26], which should be reflected in higher detection thresholds in older than young adults when postural demands are high.

## Methods

### Participants and apparatus

We recruited 23 young (24.04 ± 4.25 years old, range: 18-35; height: 171.26 ± 10.72 cm; 17♀, 6**♂**) and 20 older (64.10 ± 5.09 years old, range: 55-72; height: 170.15 ± 10.76 cm; 15♀, 5**♂**) healthy participants. Participants were free from any known neurological or musculoskeletal issues at the moment of the experiment and had normal or corrected-to-normal vision. Upon their arrival, participants signed an informed consent approved by the local ethics committee of the Justus Liebig University Giessen in accordance with the “World Medical Association Declaration of Helsinki” (2013, except for §35, pre-registration). At the end of the experiment, participants received either 8€/hour or course credits.

A custom-made vibrotactile stimulation device (Engineer Acoustics Inc., Florida, US), which will be referred to as a “tactor”, was attached to the *musculus gastrocnemius lateralis* of the participant’s right calf, a muscle primarily used for balance control [30]. This tactor was fixed at 3 cm medial of 2/3 of an imaginary line connecting the heel with the *caput fibulae*. Thus, the tactile device was fixed on the skin on top of the muscle belly. Care was taken that loose wires or clothing could not be mistaken as vibration and would not hinder the performance of the task. To manipulate the postural demands, we asked participants to either sit, stand on a rigid surface, or stand on a piece of foam, similarly to previous work [31,32]. We used a force plate (AccuSway, AMTI, Massachusetts, US) that sampled the center of pressure (COP) at 300 Hz to assess postural behaviour. In the sitting condition, participants sat on a chair with their feet on the force plate and their knees at ∼90 deg. In the standing condition, they stood directly on the force plate (∼5 cm from the ground) while in the foam condition participants stood on a balance pad (50 × 40 × 6 cm; SISSEL BalanceFit Pad, novacare GmbH, Bad Dürkheim, Germany) that was placed on top of the force plate. Participants were either barefoot or with their socks on. To reduce the fear of falling an additional experimenter stood next to the participant. For the standing conditions, participants were upright with their arms relaxed at their sides and had to fixate a circular point (18 cm diameter) on the facing wall, ∼230 cm in front of them and at a height of 185 cm.

### Procedure

Data acquisition as well as pre-processing and analysis were handled by custom-made Matlab R2021a software (The Mathworks, Massachusetts, US).

The experiment consisted of 5 phases (familiarization, training, and three experimental blocks). During familiarization, we presented 5 trials with a vibrotactile stimulus (50 ms, 250 Hz) to the participant’s leg while standing, ranging from low (peak-to-peak amplitude: 31.6 μm) to strong (peak-to-peak amplitude: 284.4 μm) intensities. This familiarization phase allowed participants to understand how the stimuli felt. Please note that this vibration is irrelevant for movement control and is used to assess tactile perception on the calf.

Afterward, a training phase followed the experimental procedure. Participants stood on the force plate for 10 trials, with a vibrotactile stimulus presented randomly between 2 and 3 seconds after trial onset (controlled by the experimenter via button press). The stimulus intensity for each trial was determined using a QUEST algorithm (Psychtoolbox Version 3.0.18), which employs a Bayesian approach to estimate psychometric function parameters using prior knowledge. We used a Weibull function with β = 3.5, as recommended for 2AFC tasks [33]. Prior mean and standard deviation were estimated from a pilot study with young participants standing with their feet closed (N=16; exposed to 50 stimuli of varying intensities). Participants responded verbally when they felt a stimulation, and an auditory cue at the end of each trial prompted them to respond if they had not done so already. An experimenter recorded the responses via button presses.

This training phase was followed by the 3 experimental blocks, each consisting of a single condition (sitting, standing, foam; Fig. 1), presented in random order. Every block started with 20 trials with the intensities of the tactile probes decided by a QUEST algorithm that had the same parameters as those described above. We then used the mean of the estimated probability density function of these 20 trials to construct a range of stimuli for that condition and participant. We used this range of stimuli to assess tactile perception with the method of constant stimuli during the experimental blocks. By first estimating a threshold with the QUEST procedure and tailoring our stimuli around that threshold we minimized the risk of presenting stimuli that were too weak or too strong for individual participants. We chose 6 stimulus intensities that were weaker and another 6 intensities that were stronger than the threshold estimated by the QUEST, resulting in a total of 13 intensities separated by a gain of 6.32 μm. When the estimated threshold by the QUEST was lower than 41.08 μm, we were unable to choose 6 unique intensities below that threshold. In such cases (17 blocks in 11 participants) every level that would correspond to an intensity of 0 or lower was handled as a level without stimulation (catch trial). The resulting range of presented stimuli can therefore be different for every condition and participant to optimize psychometric probing and fitting. We presented 6 trials for each stimulus intensity and another 22 mandatory catch (no-stimulation) trials resulting in a total of 100 trials per condition. The intensities were presented in a random order within each block. While participants were allowed to take breaks whenever they wanted, mandatory breaks of at least one minute were enforced during both standing conditions after 20 QUEST trials and after every 25^th^ trial. Similarly, to the training phase, the tactile probe was presented within 2-3 seconds after trial onset.

**Figure 1.**
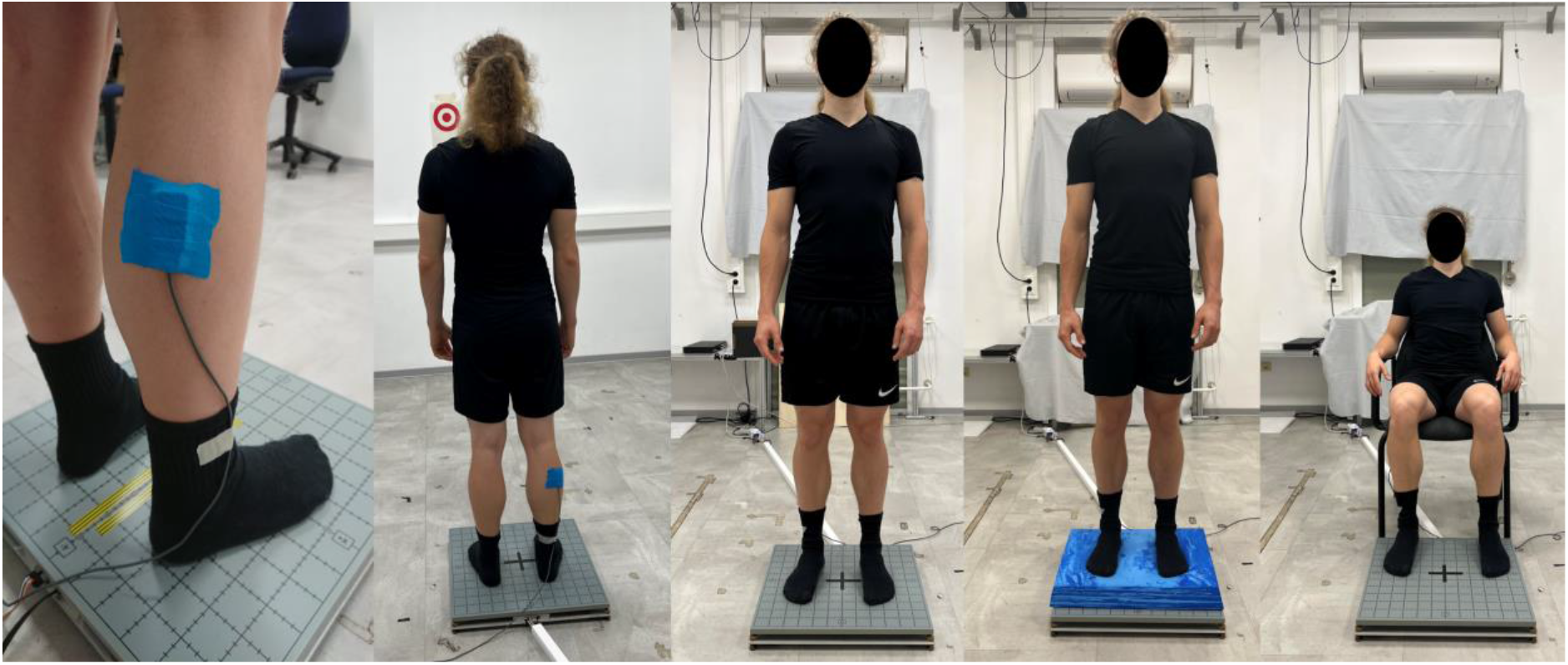
The experimental setup. Pictures of the tactor placement, the target location (red circle), the stable standing, the unstable standing, and the sitting condition (left to right).

### Data analysis

To evaluate how postural conditions affected upright stance, we analyzed COP characteristics, focusing on standing conditions as COP remains stable while sitting. We pre-processed the COP data by subtracting the mean mediolateral and anteroposterior values from each trial’s time-series data. We excluded unrealistic outliers (values above 15 cm and below -15 cm) and smoothed the data using a 100 ms moving average. Since the stimulation started the earliest 2 seconds after the trial began, we standardized the COP data to the first 2 seconds to exclude any influence of the tactile probe on posture, allowing better comparability. For each trial, we calculated the area of a 95% confidence interval ellipse (95EA) and the two-dimensional COP path length (COP_Length_), then averaged these values per condition and participant. To evaluate how tactile signals are influenced by postural conditions, we fitted the responses to tactile stimuli in each condition with a Weibull function using the Psignifit 4 toolbox [34] in Matlab. We defined the tactile detection threshold as the stimulus intensity at a 50% detection rate, adjusting for lapse and guess rates. The slope at this threshold served as a measure of perceptual sensitivity. To account for individual differences, we normalized each participant’s detection thresholds and slopes by subtracting their sitting baseline values from those obtained in the two standing conditions, resulting in two normalized threshold and slope values per participant. Three kinematic datasets of young participants were excluded due to a technical malfunction of the force plate. Since we also performed analysis solely on psychophysical data, we kept the remaining data to increase statistical power. Psychophysical data was excluded if false positive rates exceeded 20% or if the estimated detection threshold was outside the stimulated range (not counting catch trials as part of the range). This resulted in the exclusion of 7% of the total detection thresholds and slopes. Afterward, we corrected for outliers which we defined as data points outside of a 3 interquartile range. This led to an exclusion of 3.3% and 1.5% of the total kinematic data and of 3% and 1.7% of the total psychophysical data in older and young adults, respectively.

### Statistical analysis

To examine if postural demands affect upright stance in young and older adults, we conducted two separate 2 (age groups) x 2 (postural demands) repeated measures mixed ANOVAs on 95EA and COP_Length_. To confirm that aging reduces tactile perception, we compared detection thresholds and slopes during the sitting baseline between age groups using one-sided paired t-tests. We also conducted two separate 2 × 2 repeated measures mixed ANOVAs to analyze the influence of age and postural demands on normalized detection thresholds and slopes. To explore any potential relationship between the relative influence of postural demands and the relative modulation of tactile perception, we calculated for each participant (a) the change in postural sway and (b) the change in tactile perception measures (thresholds, slopes) between the two standing conditions. We then examined whether there is any relationship between the change in each of 95EA and COP_Length_ and the changes in the detection thresholds and slopes by calculating Pearson correlation coefficients.

We set our alpha to 0.05 and we report effect sizes as *η*^*2*^ following the calculations and recommendations [35]. Statistical analyses were conducted in JASP version 0.18.1 (University of Amsterdam, The Netherlands).

## Results

### Kinematics

As expected, 95EA and COP_Length_ were higher when standing on foam than on the rigid surface (*F*_*1,36*_ = 49.693, *p* < 0.001, *η*_*part*_^*2*^ = 0.580; *F*_*1,38*_ = 46.658, *p* < 0.001, *η*_*part*_^*2*^ = 0.551; Figure 2), confirming that our experimental manipulation led to the expected increase in postural sway. However, no main effect of age *(F*_*1,36*_ = 3.074, *p* = 0.088, *η*_*part*_^*2*^ = 0.079; *F*_*1,38*_ = 0.002, *p* = 0.965, *η*_*part*_^*2*^ < 0.001) or interactions *(F*_*1,36*_ = 0.003, *p* = 0.937, *η*_*part*_^*2*^ < 0.001; *F*_*1,38*_ = 0.648, *p* = 0.426, *η*_*part*_^*2*^ = 0.017) were found.

**Figure 2.**
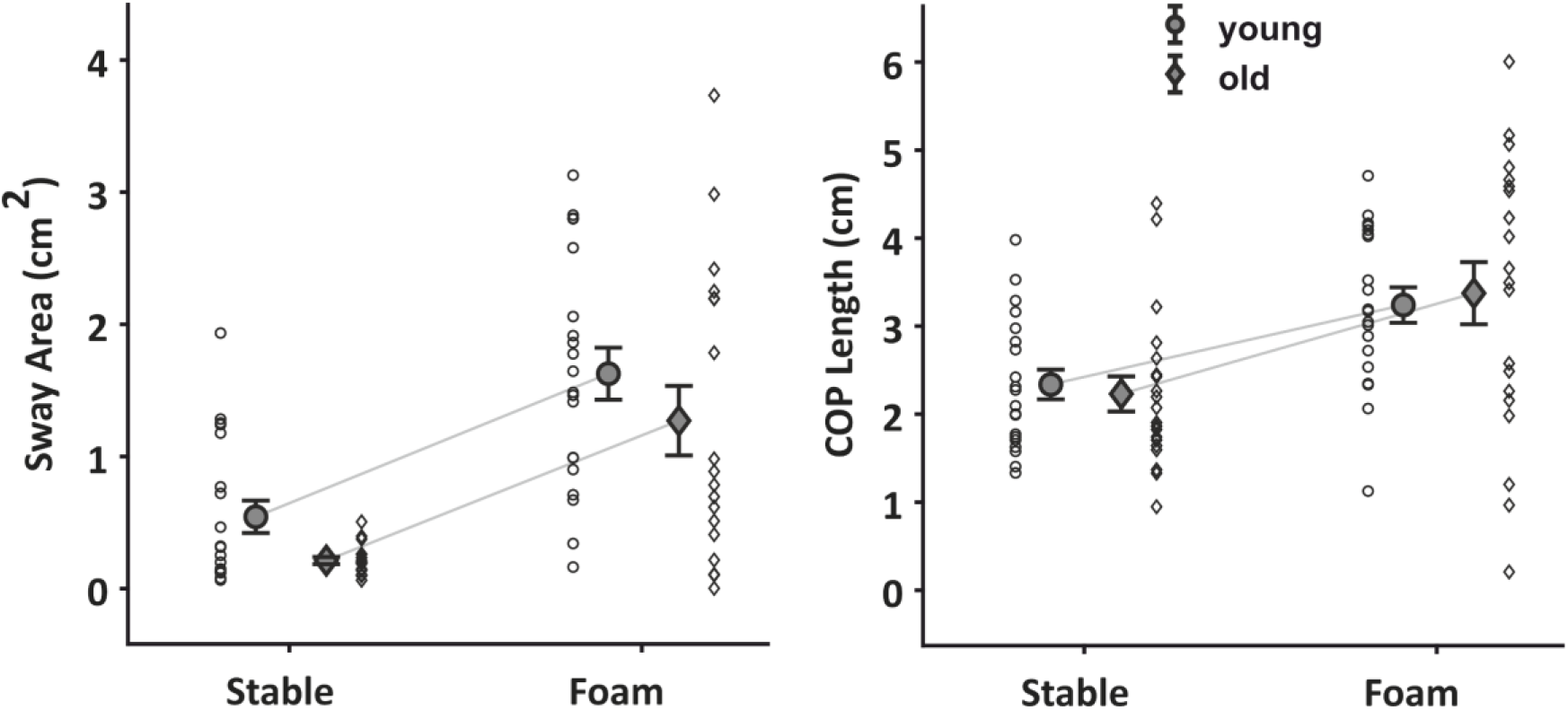
Kinematic modulation. Comparison between postural demands and age group for sway area (left) and center of pressure (COP) length (right). Means and single subject data for the young (circles) and older adults (diamonds) are depicted with standard error as error bars.

### Tactile perception

The one-sided independent t-tests showed that older adults had overall higher baseline detection thresholds than young adults (*t*_*41*_ = 1.845, *p* = 0.036, *η* = 0.073). However, there were no statistically significant differences in the slopes of the baseline psychometric functions between the two age groups (*t*_*38*_ = 0.640, *p* = 0.737, *η* = 0.010; Figure 3).

**Figure 3.**
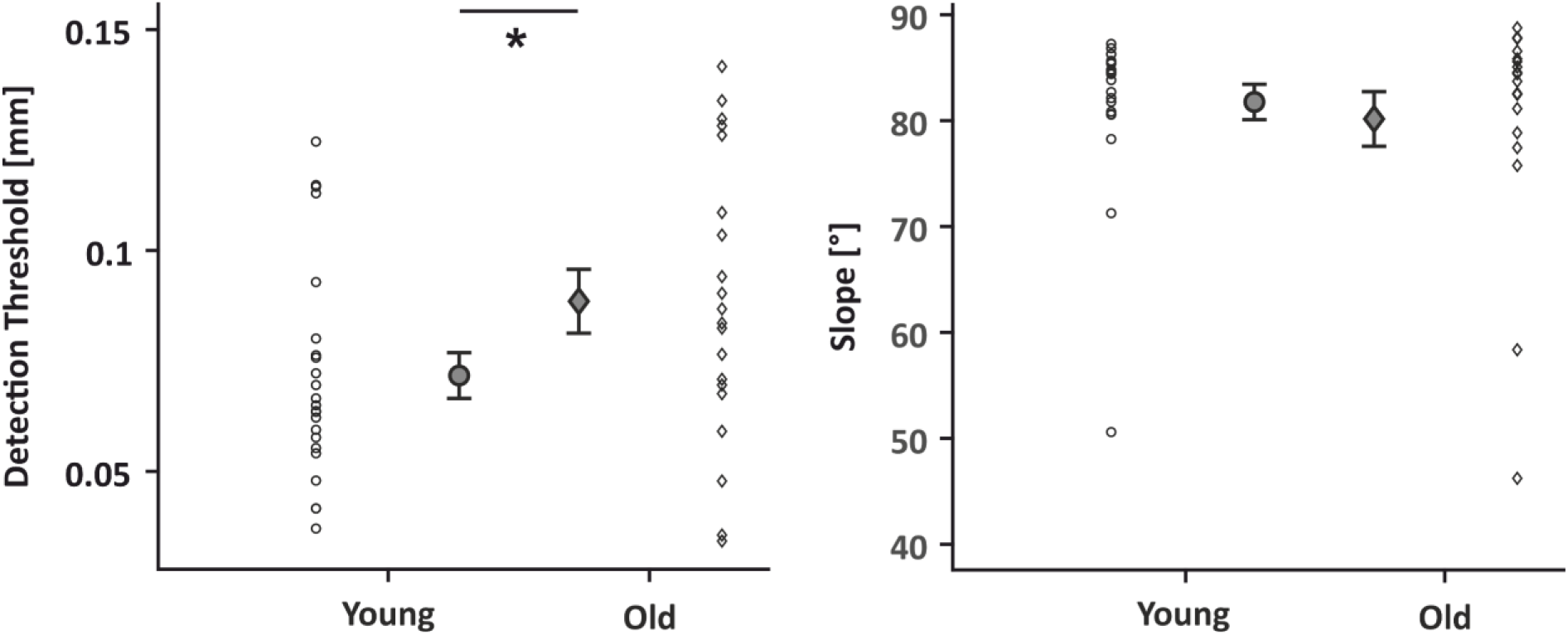
Age effects on tactile perception. Comparison between the age groups for detection thresholds (left) and slope (right) of the psychometric function in the baseline condition. Means and single subject data for the young (circles) and older adults (diamonds) are depicted with standard error as error bars. * p < 0.05

To evaluate how aging and postural demands influence tactile modulation, we ran two 2×2 ANOVAs on the normalized thresholds and slopes. Tactile thresholds were higher when standing on foam than on the rigid surface (*F*_*1,41*_ = 8.285, *p* = 0.006, *η*_*part*_^*2*^ = 0.168), but there was again no main effect of age *(F*_*1,41*_ = 0.611, *p* = 0.439, *η*_*part*_^*2*^ = 0.015) or an interaction (*F*_*1,41*_ = 0.790, *p* = 0.379, *η*_*part*_^*2*^ = 0.019; Figure 4.

**Figure 4.**
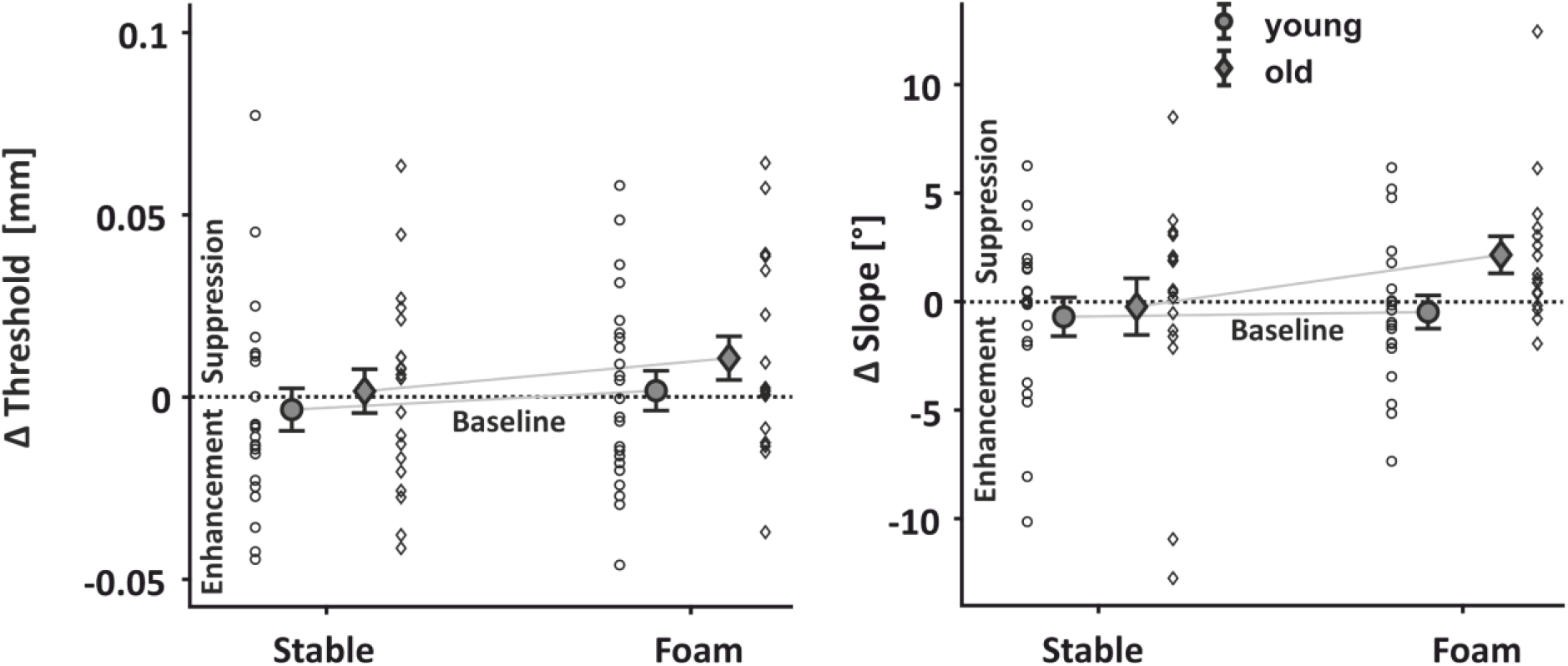
Effect of age and postural demands on tactile perception. Comparison between postural demands and age group for normalized detection thresholds (left) and normalized slopes (right). Means and single subject data for the young (circles) and older adults (diamonds) are depicted with standard error as error bars. Dotted lines represent values at sitting (baseline).

There were also no main effects or interaction for the slopes *(F*_*1,34*_ <= 2.948, *p* >= 0.095, *η*_*part*_^*2*^ *<*= 0.080).

To test for potential relationships between the elicited sway by the postural demands and the changes in tactile perception we conducted four correlations comparing the two measures for sway with the two measures for tactile perception. None of the four correlations reached significance (*r* <= 0.27, *p* >= 0.135, *η*^*2*^ <= 0.073).

## Discussion

We demonstrate increased postural sway when standing on foam than on a rigid surface, supporting previous findings [e.g. 36,21]. This confirms that the underlying postural demands imposed by standing on foam can modulate posture. As expected ([17,26], we found that aging leads to overall poorer tactile perception. Importantly, tactile perception became worse with increasing postural demands, but this modulation was independent of age.

Tactile sensitivity was poorer when standing on an unstable than a stable surface. This fits previous work showing poorer tactile sensitivity when postural demands increase, such as when standing compared to sitting [37]. Thus, humans seem to perceive and incorporate less tactile information when posture is challenged. One possible explanation for this effect might be that increased task demands led to worse tactile sensitivity, as shown for arm movements [15, but see also 16]. Another possibility is that afferent signals from the leg were stronger when standing on foam than on a rigid surface, for instance due to the larger sway or associated muscle activity, and that these afferent signals may have masked the tactile probe we used to assess tactile sensitivity. Previous studies suggest that backward masking due to a high afferent signal may at least partly explain tactile suppression [11], though other studies did not find such an influence [e.g. 38,39]. While the precise mechanisms are yet to be determined, we see a clear decrease in tactile perception with postural demand.

We do not have any evidence that postural demands influence tactile sensitivity differently in young and old humans. This might be because aging did not influence postural sway in either direction. There is work showing that aging leads to increased postural sway [22], although there is some evidence showing the opposite effect [23]. One difference between previous studies and our one is that we measured postural sway during a limited duration of 2 seconds, whereas previous studies on the role of aging in postural control assessed posture during longer time periods (> 10 seconds) [ e.g. 20,22,23]. Here, we used shorter durations to keep our experiment within a reasonable duration for the older participants given the high number of trials needed for robust psychophysical measures. Thus, the absence of a differential effect of postural demands on tactile modulation for young and older participants might stem from the postural demands not causing different postural responses between age groups.

We also explored whether the increased postural sway in the foam compared to the standing condition could explain the associated tactile modulation, as the movement itself may lead to tactile suppression [40]. However, the increase in detection thresholds and slopes between the two standing conditions was not correlated with the associated increase in postural sway leaving open whether and how psychophysical and postural measures relate to each other.

In sum, tactile perception was hampered in older compared to young participants and tactile perception was reduced with higher postural demands. However, we have no evidence that aging comes with stronger impairment in tactile perception when postural demands are high. This suggest that postural demands challenge healthy agers as young adults, probably leading to a down-weighting of tactile feedback processing.

## Data availability

Behavioral and psychophysical data are publicly available after acceptance at https://osf.io/jwghy/

## Author contributions

Conceptualization: F.W., K.F., D.V.

Data curation: F.W.

Formal analysis: F.W.

Funding acquisition: K.F., D.V.

Investigation: F.W.

Methodology: F.W., K.F., D.V.

Project administration: F.W., K.F., D.V.

Resources: F.W., K.F., D.V.

Software: F.W.

Supervision: K.F., D.V.

Validation: F.W., K.F., D.V.

Visualization: F.W.

Roles/Writing - original draft: F.W.

and Writing - review & editing: F.W., K.F., D.V.

## Acknowledgments

This work was supported by the Collaborative Research Center SFB/TRR 135, project A4, under grant agreement 222641018 funded by the German Research Foundation (Deutsche Forschungsgemeinschaft), by “The Adaptive Mind” funded by the Hessian Ministry for Higher Education, Research, Science and the Arts, and by the German Research Foundation under grant agreement VO 2542/1-1. We would like to thank Kani Gaff and Viktoria Wild for their assistance in data collection.

## References

[1] Chiba R, Takakusaki K, Ota J, Yozu A, Haga N. Human upright posture control models based on multisensory inputs; in fast and slow dynamics. Neuroscience Research 2016;104:96–104. 10.1016/j.neures.2015.12.002.

[2] Peterka RJ. Sensory integration for human balance control. Handbook of Clinical Neurology, vol. 159, Elsevier; 2018, p. 27–42. 10.1016/B978-0-444-63916-5.00002-1.

[3] Wolpert DM, Ghahramani Z. Computational principles of movement neuroscience. Nat Neurosci 2000;3:1212–7. 10.1038/81497.

[4] Buckingham G, Carey DP, Colino FL, deGrosbois J, Binsted G. Gating of vibrotactile detection during visually guided bimanual reaches. Exp Brain Res 2010;201:411–9. 10.1007/s00221-009-2050-8.

[5] Voudouris D, Broda MD, Fiehler K. Anticipatory grasping control modulates somatosensory perception. Journal of Vision 2019;19:1–10. 10.1167/19.5.4.

[6] Mildren RL, Strzalkowski NDJ, Bent LR. Foot sole skin vibration perceptual thresholds are elevated in a standing posture compared to sitting. Gait & Posture 2016;43:87–92. 10.1016/j.gaitpost.2015.10.027.

[7] Fuehrer E, Voudouris D, Lezkan A, Drewing K, Fiehler K. Tactile suppression stems from specific sensorimotor predictions. Proc Natl Acad Sci USA 2022;119:e2118445119. 10.1073/pnas.2118445119.

[8] Voss M, Ingram JN, Haggard P, Wolpert DM. Sensorimotor attenuation by central motor command signals in the absence of movement. Nat Neurosci 2006;9:26–7. 10.1038/nn1592.

[9] Voss M, Ingram JN, Wolpert DM, Haggard P. Mere expectation to move causes attenuation of sensory signals. PLoS One 2008;3:e2866. 10.1371/journal.pone.0002866.

[10] Williams SR, Chapman CE. Time Course and Magnitude of Movement-Related Gating of Tactile Detection in Humans. III. Effect of Motor Tasks. Journal of Neurophysiology 2002;88:1968–79. 10.1152/jn.2002.88.4.1968.

[11] Chapman CE, Beauchamp E. Differential Controls Over Tactile Detection in Humans by Motor Commands and Peripheral Reafference. Journal of Neurophysiology 2006;96:1664–75. 10.1152/jn.00214.2006.

[12] Voudouris D, Fiehler K. Dynamic temporal modulation of somatosensory processing during reaching. Sci Rep 2021;11:1928:1–12. 10.1038/s41598-021-81156-0.

[13] Voudouris D, Fiehler K. The role of grasping demands on tactile suppression. Human Movement Science 2022;83:102957. 10.1016/j.humov.2022.102957.

[14] Mouchnino L, Fontan A, Tandonnet C, Perrier J, Saradjian AH, Blouin J, et al. Facilitation of cutaneous inputs during the planning phase of gait initiation. Journal of Neurophysiology 2015;114:301–8. 10.1152/jn.00668.2014.

[15] McManus M, Schütz I, Voudouris D, Fiehler K. How visuomotor predictability and task demands affect tactile sensitivity on a moving limb during object interaction in a virtual environment. R Soc Open Sci 2023;10:1–22. 10.1098/rsos.231259.

[16] Gertz H, Fiehler K, Voudouris D. The role of visual processing on tactile suppression. PLoS ONE 2018;13:e0195396. 10.1371/journal.pone.0195396.

[17] McIntyre S, Nagi SS, McGlone F, Olausson H. The Effects of Ageing on Tactile Function in Humans. Neuroscience 2021;464:53–8. 10.1016/j.neuroscience.2021.02.015.

[18] Seidler RD, Bernard JA, Burutolu TB, Fling BW, Gordon MT, Gwin JT, et al. Motor control and aging: Links to age-related brain structural, functional, and biochemical effects. Neuroscience & Biobehavioral Reviews 2010;34:721–33. 10.1016/j.neubiorev.2009.10.005.

[19] Patel M, Fransson PA, Lush D, Gomez S. The effect of foam surface properties on postural stability assessment while standing. Gait & Posture 2008;28:649–56. 10.1016/j.gaitpost.2008.04.018.

[20] Lion A, Spada RS, Bosser G, Gauchard GC, Anello G, Bosco P, et al. “Postural first” principle when balance is challenged in elderly people. International Journal of Neuroscience 2014;124:558–66. 10.3109/00207454.2013.864288.

[21] Wallace KM, Brown MR, Pannell WC, Daniels JB, Moore JC, McInnis AK, et al. Impact of Differing Instability Devices on Postural Sway Parameters. Applied Sciences 2024;14:3029. 10.3390/app14073029.

[22] Degani AM, Leonard CT, Danna-dos-Santos A. The effects of early stages of aging on postural sway: A multiple domain balance assessment using a force platform. Journal of Biomechanics 2017;64:8–15. 10.1016/j.jbiomech.2017.08.029.

[23] Šarabon N, Kozinc Ž, Marković G. Effects of age, sex and task on postural sway during quiet stance. Gait & Posture 2022;92:60–4. 10.1016/j.gaitpost.2021.11.020.

[24] Kanekar N, Aruin AS. The effect of aging on anticipatory postural control. Exp Brain Res 2014;232:1127–36. 10.1007/s00221-014-3822-3.

[25] Hatzitaki V, Voudouris D, Nikodelis T, Amiridis IG. Visual feedback training improves postural adjustments associated with moving obstacle avoidance in elderly women. Gait & Posture 2009;29:296–9. 10.1016/j.gaitpost.2008.09.011.

[26] Klever L, Voudouris D, Fiehler K, Billino J. Age effects on sensorimotor predictions: What drives increased tactile suppression during reaching? Journal of Vision 2019;19:1–17. 10.1167/19.9.9.

[27] Wolpe N, Ingram JN, Tsvetanov KA, Geerligs L, Kievit RA, Henson RN, et al. Ageing increases reliance on sensorimotor prediction through structural and functional differences in frontostriatal circuits. Nat Commun 2016;7:13034. 10.1038/ncomms13034.

[28] Timar L, Job X, Orban De Xivry J-J, Kilteni K. Aging exerts a limited influence on the perception of self-generated and externally generated touch. Journal of Neurophysiology 2023;130:871–82. 10.1152/jn.00145.2023.

[29] World Medical Association Declaration of Helsinki: Ethical Principles for Medical Research Involving Human Subjects. JAMA 2013;310:2191. 10.1001/jama.2013.281053.

[30] Gatev P, Thomas S, Kepple T, Hallett M. Feedforward ankle strategy of balance during quiet stance in adults. The Journal of Physiology 1999;514:915–28. 10.1111/j.1469-7793.1999.915ad.x.

[31] Mademli L, Mavridi D, Bohm S, Patikas DA, Santuz A, Arampatzis A. Standing on unstable surface challenges postural control of tracking tasks and modulates neuromuscular adjustments specific to task complexity. Sci Rep 2021;11:6122. 10.1038/s41598-021-84899-y.

[32] Taneda K, Mani H, Kato N, Komizunai S, Ishikawa K, Maruya T, et al. Effects of simulated peripheral visual field loss on the static postural control in young healthy adults. Gait & Posture 2021;86:233–9. 10.1016/j.gaitpost.2021.03.011.

[33] Watson AB, Pelli DG. Quest: A Bayesian adaptive psychometric method. Perception & Psychophysics 1983;33:113–20. 10.3758/BF03202828.

[34] Schütt HH, Harmeling S, Macke JH, Wichmann FA. Painfree and accurate Bayesian estimation of psychometric functions for (potentially) overdispersed data. Vision Res 2016;122:105–23. 10.1016/j.visres.2016.02.002.

[35] Correll J, Mellinger C, McClelland GH, Judd CM. Avoid Cohen’s ‘Small’, ‘Medium’, and ‘Large’ for Power Analysis. Trends in Cognitive Sciences 2020;24:200–7. 10.1016/j.tics.2019.12.009.

[36] Liu B, Leng Y, Zhou R, Liu J, Liu D, Liu J, et al. Foam pad of appropriate thickness can improve diagnostic value of foam posturography in detecting postural instability. Acta Oto-Laryngologica 2018;138:351–6. 10.1080/00016489.2017.1393842.

[37] Wachsmann FD, Fiehler K, Voudouris D. Temporal modulation of tactile perception during balance control. BioRxiv 2024. 10.1101/2023.11.02.565260.

[38] Arikan BE, Voudouris D, Straube B, Fiehler K. Distinct role of central predictive mechanisms in tactile perception. OSF 2023. 10.31234/osf.io/rqs9p.

[39] Broda MD, Fiehler K, Voudouris D. The influence of afferent input on somatosensory suppression during grasping. Sci Rep 2020;10:18692:1–11. 10.1038/s41598-020-75610-8.

[40] Chapman CE, Bushnell MC, Miron D, Duncan GH, Lund JP. Sensory perception during movement in man. Exp Brain Res 1987;68. 10.1007/BF00249795.

